# Deconstruction of the retrosplenial granular cortex for social behavior in the mouse model of fragile X syndrome

**DOI:** 10.1101/2021.01.24.428008

**Authors:** Hui-Fang Shang, Ruonan Cai, Hao Sun, Tao Sheng, Yan-Na Lian, Li Liu, Wei Chen, Lixia Gao, Han Xu, Chen Zhang, Jian-Hong Luo, Xinjian Li, Xiang-Yao Li

## Abstract

Deficits in fragile X mental retardation 1 protein lead to fragile X syndrome (FXS) with mental retardation and social activity disorder. Until now, the neuronal circuits that mediate the social impairments of FXS were mostly unclear. Accidently, we found fewer c-fos expression in RSG of KO than WT mice after social behavior test. Inactivation of RSG neurons decreased social novelty but not the sociability of naive mice. Interestingly, although the RSG neurons of KO mice had higher background activity, fewer social contact-related Ca^2+^ neurons were observed during social interaction test *via* one-photon Ca^2+^ imaging in freely-behaving mice. Strikingly, enhancing the activity of RSG neurons rescued the abnormal social novelty in KO mice. Further studies proved that the innervations from the subiculum and ACC to RSG contributes to the social behavior. Take together, we found that abnormal activity in the retrosplenial granular cortex (RSG) led to social novelty deficits in *Fmr1*-knockout (KO) mice. Moreover, selective manipulation of RSG neurons may be an effective strategy to treat the social deficits in FXS.

**One Sentence Summary:** Deletion of FMRP leads to lower social-related neuronal activity in the RSG; this causes social novelty deficits in *Fmr1*-KO mice.

## INTRODUCTION

Social behaviors include a serial of complex fundamental behaviors that are essential for life in society. Previous studies have indicated appropriate social activity depends on the appropriate neural activities in different brain regions, including the medial prefrontal cortex (mPFC), hippocampus, amygdala, midbrain, and nucleus accumbens (NAc) (Chen and Hong, 2018; Fernández et al., 2018). Genetic and environmental factors may change neural activity in social-related brain regions, impairs the processing of social-related sensory information, leading to some psychiatric disorders (Barak and Feng, 2016; Chen and Hong, 2018), such as autism spectrum disorder, schizophrenia, and depression. Although it has been widely studied, the neural mechanism of social behavior is far more unclear. Thus, studies on the neural circuits that mediate abnormal social interactions would help understand social cognition mechanisms and provide opportunities to treat the prevailing social disorders (Barak and Feng, 2016).

During social interaction, the subject needs to integrate multiple sensory inputs, decide, and perform behaviors (Kingsbury and Hong, 2020), which indicates that most social-related brain regions integrate different information and have dense connections to other brain regions. Anatomically, the retrosplenial cortex(RSC) reciprocally connected with many brain regions, including the hippocampus, thalamus, visual cortex, auditory cortex, motor cortex, and prefrontal cortex (Jiang et al., 2018; Todd et al., 2016). Most of them are key brain regions for social behavior, which is indicated by RSC’s function, such as spatial navigation (11), emotion, learning and memory, and high cognitive functions: imagination. These results imply that RSC may integrate sensory-motor activity during social behavior. A recent study has linked the neuronal activities in layer V of RSCs to social interaction (Vesuna et al., 2020). However, few studies have been performed on its role in social activities, especially its connection to FXS social disorder.

Deletion of the fragile X mental retardation gene (*Fmr1*) leads to fragile X syndrome (FXS), the most common inherited developmental disorder (Hagerman et al., 2017; Melancia and Trezza, 2018). Fragile X mental retardation protein (FMRP) is an RNA-binding protein that mainly translocate mRNA and regulates the translation of various synapse-related proteins (Hagerman et al., 2017; Kwan et al., 2012). Deficits in FMRP affect excitatory and inhibitory synaptic transmission (Gao et al., 2018) and synaptic plasticity (Hagerman et al., 2017), leading to functional changes at the level of neuronal circuits in the olfactory system (Daroles et al., 2016), brain stem (Ruby et al., 2015), and cerebral cortex (Kwan et al., 2012). The core cognitive features of FXS include mental retardation and social disability (Melancia and Trezza, 2018). Like humans, mice with FMRP deficits (*Fmr1* mutant) show similar social problems, including social withdrawal, anxiety, and social cognition deficits. However, the neural mechanism underlying social deficits in FXS is mainly unknown.

Here, we found a decrease of c-fos-positive neurons in the RSG of *Fmr1*-knockout (KO) mice after social interaction. We confirmed the involvement of RSG in social novelty but not sociability using optogenetic and chemogenetic approaches. Moreover, *in vivo* Ca^2+^ imaging revealed that insufficient neuronal activity occurred in the RSG in sociability and social novelty stages in *Fmr1*-KO mice. Furthermore, inhibiting the projection from subiculum and ACC to RSG impaired social novelty. Thus, we provide the strong evidence that the RSG is involved in social cognition.

### Materials and methods

#### Experiment model and subject details

Adult male C57B L/6 mice (aged 8 weeks at the start of experimental procedures) were purchased from Shanghai SLAC Laboratory Animal Co… *Fmr1* KO mice (FVB.129P2(B6)-*Fmr1*^tm1Cgr^/J, stock number 003024) on an FVB genetic background were purchased from the Jackson Laboratory. Mice were bred and maintained in the experimental animal center of Zhejiang University. *Fmr1* KO mice were generated by crossing heterozygous females with heterozygous male mice. The offsprings were genotyped by PCR using mouse tail-tip DNA and common primers (5′- CTT CTG GCA CCT CCA GCT T-3′), WT primers (5′- TGT GAT AGA ATA TGC AGC ATG TGA-3′) for 131 bp and mutant allele-specific primers (5′- CAC GAG ACT AGT GAG ACG TG-3′) for 400 bp. The PCR products were visualized with ethidium bromide staining. For experiments, mice (20 - 35 g) were housed 4 or 5 per cage at constant room temperature (25 ± 1°C) and relative humidity (60 ± 5%) under a 12-hr light/dark schedule (lights on 07:00-19:00); food and water were available *ad libitum*. For all behavioral tests, the mice were allowed to adapt to laboratory conditions for about one week and habituate to the testing situation for at least 15 min before experiments. The Animal Care and Use Committee of Zhejiang University approved all of the mouse protocols.

## METHOD DETAILS

### Stereotaxic Virus Injection

Stereotaxic Injections of AAVs were performed as described (Wang et al., 2020). Briefly, mice (6-8 weeks old) were anesthetized with isoflurane (induction 4%, maintenance 1%), and the scalp was shaved and cleaned with iodine (Triadine). The mouse’s head was fixed into a stereotaxic adapter mounted on a stereotaxic frame (Kopf model 962), and lubricant (Artificial Tears) was applied to the eyes. An incision was made over the skull, and the surface was exposed. Two small holes (1 hole per hemisphere) were drilled above the RSC (AP:-2.06mm, ML: +0.05mm, VD:-1.05mm), and the dura was gently reflected. The virus was infused at a rate of 20 nl per min.

Following infusion, the needle was kept at the injection site for 10 min and then slowly withdrawn. The total amount of virus (10^12, 600 nl/animal) infused was based on its titer. Two more extra virus injection sites were introduced at the interested brain region to study the neural circuit that related to RSC. We injected RSC with pAOV2/Retro-CamkⅡα-eYFP-2A-Cre virus and 7 days later injected cg1/SUB with AAV-Ef1α-DIO-eNpHR3.0-mCherry virus. Then optic fiber was implanted in Cg1/SUB to specifically inhibit retrograde neural projections from RSG.

### Cannula Implantation

The cannula implantation and microinjection were performed as described previously (Wang et al., 2020). In brief, mice were anesthetized with isoflurane inhalation (1-3%, as needed; R510-22, RWD, Shenzhen, China) in 100% oxygen at 0.5 L/min delivered by the facemask. The scalp was shaved and cleaned with iodine (Caoshanhu, Nanchang, China) and alcohol. The head of each mouse was fixed into a stereotaxic adapter mounted on a stereotaxic frame (68025, RWD, Shenzhen, China), and ointment (Cisen, Jining, China) was applied to the eyes. An incision was made over the skull, and the surface was exposed. Two small holes were drilled above the RSC, and the dura was gently reflected. Guide cannulas were placed −2.1 mm anterior to bregma, 0.5 mm lateral to the midline, and 1.05 mm ventral to the surface of the skull. For microinjection, the mice were restrained in a plastic cone (68025, RWD), and a small hole was cut in the plastic overlying the microinjection guides. The dummy cannulas were removed, and the microinjection cannula was inserted into the guide. A 30-gauge injection cannula was placed 0.7 mm below the end of the guide. Vehicle or D (-)-2-amino-5-phosphonopentanoic acid; 0.05 mg/ml in physiological saline (R&D, CAS No: 76326-31-3) was delivered bilaterally at 0.5 μL/min using a syringe driven by an infusion pump (ALC-IP600, Alcott, Shanghai, China). After delivery to each side of the brain, the injection cannula was left in place for 2 min to minimize back-flow up the guide. The cannula was then retracted and inserted into the opposite side of the brain. Fifteen minutes after microinjection, the mice were used to do social interaction tests and other behavioral tests.

### Three-Chamber Sociability Test

The experiment was performed as described previously (Chung et al., 2015; Lo et al., 2016); 40 cm width × 20 cm height × 26 cm depth with a 12-cm-wide center chamber and 14-cm-wide side chambers. Both side chambers contained a plastic cage in the corner, with a plastic cup with a weight on it, to prevent the subject mouse from climbing. The assay consisted of four sessions. The first session began with 10-min habituation in the center chamber, followed by the second 10 min session, where the subject mouse could freely explore all three chambers. The mouse was then gently confined in the center chamber while a novel object and a wild-type stranger mouse, stranger 1, was placed in the two plastic cages. The subject mouse was then allowed to explore all three chambers for 10 min freely. Before the last session, the subject mouse was again gently guided to the center chamber while the object was replaced with a wild-type mouse, stranger 2. The subject mouse again freely explored all three chambers for 10 min. All stranger mice were males of the same age and previously habituated to the plastic cage during the previous day (30 min). The positions of object and stranger 1 were alternated between tests to prevent side preference.

### Stereotyped behavior

Continuous repetition of one or a few acts is a characteristic feature of stereotyped behavior that occurs frequently in patients with Autism Spectrum Disorders. We conducted the following quantitative, observational method for the assessment of stereotyped behavior in *Fmr1* WT or KO mice. Mice were habituated to the transparent test cages for 10min. The testing room should remain at low level white noise to screen out extraneous sounds for ongoing stereotypy can be easily disrupted by unexpected acoustic stimuli. Then recorded 10 min via video monitor for further stereotyped behavior analysis. Then animals are observed and rated via Any-maze by an experimenter who is blind to genotype. Typical stereotyped behavior as grooming, rearing, digging was scored and rated.

### Open Field Test

The experiment was performed as described previously (Wang et al., 2020). White plastic boxes were used as open-field chambers (dimensions: 45 × 45 × 45 cm^3^). Mice were individually placed in the center of a chamber and allowed to explore for 15 min freely. The locomotor and exploratory behaviors were recorded with ANY-maze software (Stoelting, Wood Dale, IL 60191, USA). The total distance traveled was used to evaluate locomotor activity.

### Elevated Plus Maze

The experiment was performed as described previously (Wang et al., 2020). The elevated plus-maze comprised two open arms (30×5 cm) without walls and two enclosed arms (30 × 5 cm) with 15-cm-high walls on each side. The arms were 60 cm above the ground. Again, all mice were individually tested in one 5-min session each. The maze was cleaned thoroughly with 75% ethanol before each test. Movements were tracked and analyzed off-line using ANY-maze software (Stoelting, Wood Dale, IL 60191, USA).

### In vivo one-photon calcium imaging with miniscope

Frm1 and littermate were used for calcium imaging in these experiments. Surgery and imaging were following previously published papers (Li et al., 2017; Liu et al., 2018). Adult mice were anesthetized with sodium pentobarbital (1%, 40mg/kg, i.p.) and treated with dexamethasone (0.2 mg/kg, s.c) to prevent brain swelling and inflammation. A piece of skull (3 mm in diameter) above the RSC was removed after high-speed dental drilling during the procedures. AAV-hSyn-GCaMP6s-WPRE (titer: 10^13, 0.6 ul/) was perfused to the RSC, then 0.6ul of the virus was added to the surface of the brain and covered by a glass coverslip (diameter, 3mm, thickness: 100 um). The scalp was closed by suturing after surgery. Three weeks after injection, a baseplate of a miniaturized integrated fluorescent microscope (Inscopix, #1050-002192) was fixed on top of glass coverslip in well-labeling animals.

Calcium imaging was performed and analyzed following our previously published papers (Li et al., 2017; Liu et al., 2018). 3-4 days post baseplate surgery, animals were habituated to the microscope’s attachment and environment for two days. Calcium imaging was then performed in a three-chamber social test apparatus following our previously published papers (Inscopix; LED power: 0.6 - 1.0 mW; camera resolution: 1,440 × 1,080 pixels). Mouse behavior was recorded with a video camera synchronized with calcium imaging using the trigger-out signal from nVista HD. Calcium imaging videos were analyzed and neuronal signals were detected using Imaging Data Processing (Inscopix) and custom-written scripts in Matlab following published algorithms (PCA-ICA).

For each neuron, each detected calcium activity’s amplitude was normalized by the standard deviation of the whole calcium trace. To visualize the detected neuron’s activity patterns during stimulation, each RSC neuron’s active event traces were aligned when the animal was directly in contact with restrained animals or an empty cup. The resulting traces from all RSC neurons were sorted by their peak activation time during the window and displayed in temporal raster plots. To calculate the activated neurons, we quantity the neurons that showed higher activity during directed social interaction.

### Immunostaining

Mice were anesthetized with 1% pentobarbital sodium and transcardially perfused with 0.1 M phosphate-buffered saline (PBS) followed by 4% paraformaldehyde (PFA) in PBS. Brains were post-fixed overnight in PFA at 4°C. Each brain was then dissected, and further fixed in 4% PFA for an additional 24 h, then transferred to 15% sucrose in PBS followed by 30% sucrose until saturated. The brain was embedded in Tissue-Tek OCT compound, frozen in liquid nitrogen, and stored at −80 °C before being cut into 25-μm coronal sections in a cryostat at –20°C (CM3050S, Leica). Free-floating sections were washed with PBS. For immunostaining, sections were incubated with blocking buffer (5% normal goat serum and 0.3% Triton X-100 in PBS) for one h at room temperature and incubated with primary antibody (c-fos, 1: 1000, Santa Cruz, CA, USA) overnight at 4°C. Sections were washed in PBS and then incubated with the appropriate secondary antibody (goat anti-rabbit 488; 1937195, Life Technologies) for two h at room temperature. Sections were rewashed for 3×10 min in PBS. After washing, they were mounted on coverslips using Fluoroshield mounting medium with DAPI (ab104139, Abcam, Cambridge, UK) for image collection. The sections were used for imaging to check out proper viral injection sites. Confocal images were captured under a 109 or 209 objective lens (Olympus FV-1200).

### Immunohistochemistry and cell count

For c-fos positive cells’ qualification, we sampled every 30-μm section from the whole brain of *Fmr1* knockout and wild-type mice. Brain regions were outlined and compared according to the reference atlas (the Allen Mouse Brain Atlas).

### Whole-cell Patch-clamp Recording

Coronal brain slices were prepared at 300 μm on a vibrating Leica VT1200S microtome using standard procedures. Slices were transferred to a submerged recovery chamber with oxygenated (95% O2 and 5% CO2) artificial cerebrospinal fluid (ACSF) containing (in mM) 124 NaCl, 2.5 KCl, 2 CaCl2, 1 MgSO4, 25 NaHCO3, 1 NaH2PO4, and 10 glucose at room temperature for at least 1 h.The recording micropipettes (3-5 MΩ) were filled with a solution containing (in mM) 124 K-gluconate, 5 NaCl, 1 MgCl2, 0.2 EGTA, 10 HEPES, 2 MgATP, 0.1 Na3GTP, and 10 phosphocreatine disodium (adjusted to pH 7.2 with KOH). Experiments were performed in a recording chamber on the stage of a microscope equipped with infrared differential interference contrast optics for visualization. The initial access resistance (15–30 MΩ) was monitored throughout experiments. Data were discarded if the access resistance changed >15% during an experiment. Data were filtered at 1 kHz and digitized at 10 kHz.

## QUANTIFICATION AND STATISTICAL ANALYSIS

Off-line analysis of whole-cell patch-clamp data was performed using Clampfit 10. GraphPad Prism 8.0 was used to plot and fit the data. Statistical comparisons were made using Student’s *t*-test, one-way ANOVA or two-way RM ANOVA (Tukey’s test, Bonferroni’s test, or Sidak’s test was used for *post hoc* comparison) or the Kruskal-Wallis test (Dunn’s Multiple Comparison Test was used for *post hoc* comparison). All data are presented as the mean ± S.E.M. In all cases, *P*<0.05 was considered statistically significant.

## RESULTS

### Involvement of the RSG in Social Interaction

Patients with FXS and mouse models of *Fmr1* mutations exhibit social disorder (Loesch et al., 2007; Spencer et al., 2005, 2008). To confirm this, we re-examined the social-related activity in *Fmr1*-KO mice using a three-chamber social interaction task (**Fig. 1A**) in which mice explored, in three sequential 10-min sessions, chambers that 1) contained two identical empty wire cup (habitation); 2) contained one familiar mouse (S1) and one empty cup (sociability); 3) contained one familiar mouse (S1) and one novelty mouse (S2, social novelty). In the sociability test, the *Fmr1*-KO mice behaved similarly to the wild-type (WT) mice, in that they spent more time exploring the unfamiliar animal than the empty cage (**Fig. 1B & C**). In the social novelty test, the WT mice spent a long time sniffing the unfamiliar mouse, while the KO mice spent the same time with both the strange and the familiar mouse (**Fig. 1B & D**). Like previous studies (Spencer et al., 2008), the absence of FMRP specifically impairs animal’s social interaction with strange animals, but not to the familiar one, which is a similar symptom of autism spectrum disorder. However, we did not observe any changes in the stereotyped behaviors of *Fmr1* KO mice, such as grooming, rearing, and digging (***fig. S1A***). Furthermore, the *Fmr1* KO mice showed standard short-term object memory (***fig. S1B***), which indicates that they have social-specific defecit compare with WT mice. So *Fmr1* KO mice showed typical social novelty deficits.

**Figure 1.**
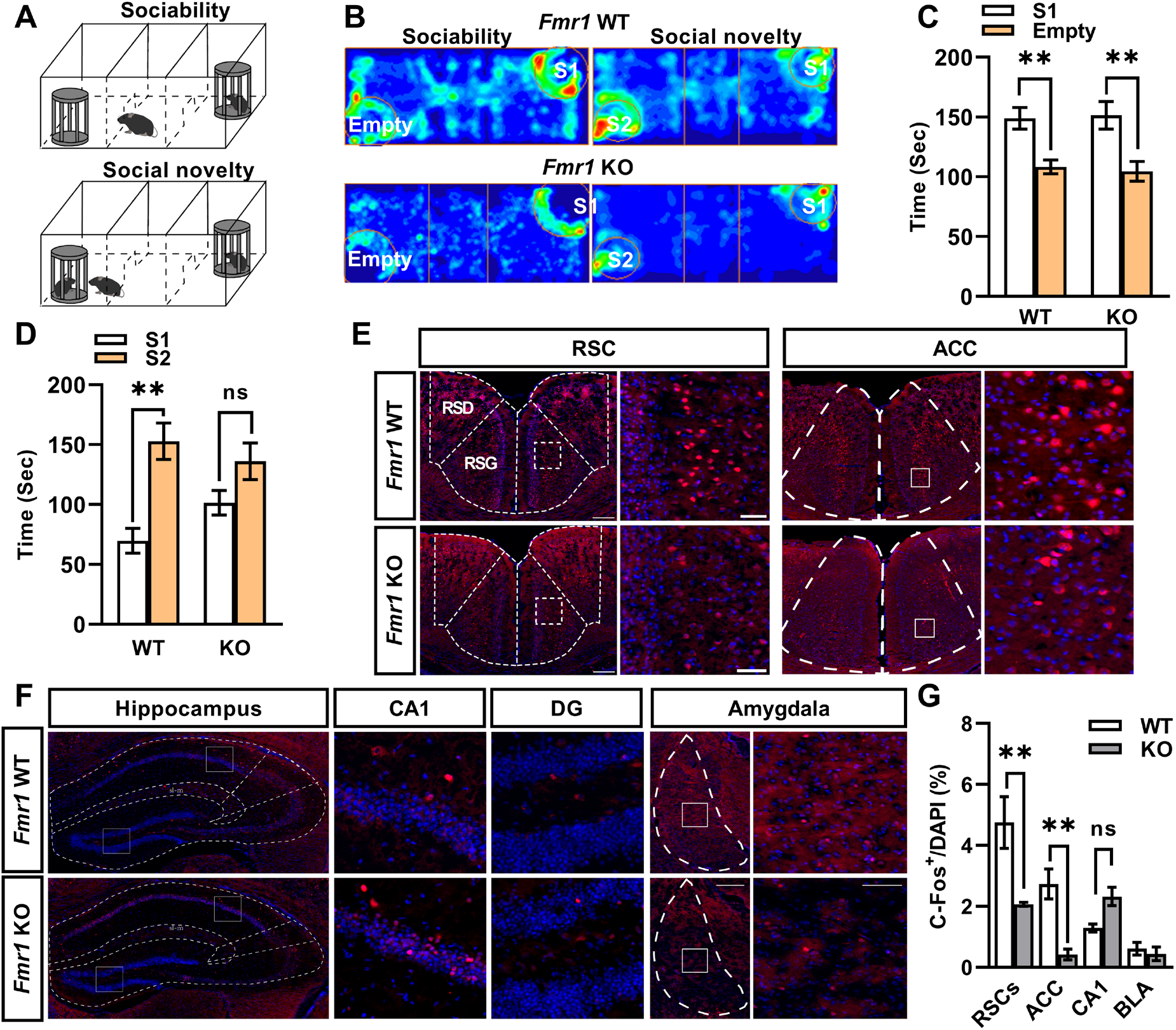
*Fmr1* KO mice showed social novelty defecit. **A**, Schematic of the social behavior test, Upper: sociability test. Lower: social novelty test. **B**, Representative mouse tracks heatmap during the sociability and social novelty tests. WT, wild-type; KO, knockout; S1, unfamiliar no.1; S2, unfamiliar no.2. **C**, WT (n = 12) and Fmr1-KO (n = 11) both prefer the S1 chamber (WT, *P* <0.01; KO, *P* <0.01) with no significant difference in interaction time (two-way ANOVA interaction, *F*_*(1,21)*_ = 26.13) during the sociability test stage. **D**, Only WT mice prefer the novel over the familiar animal (WT, *P* <0.01; KO, *P* = 0.10). **E-G**, WT (n = 3) mice, but not KO (n = 3) mice, during the social novelty test, show robust c-Fos expression in the retrosplenial granular cortex (**E**, left) and anterior cingulate cortex (**F**, right), but not in the hippocampus (**F**, left), CA1 (**F**, middle left), dentate gyrus (**F**, middle right), and amygdala (**F**, right). c-Fos-positive cells are increased in the RSG (*P* <0.01) and ACC (*P* <0.01) (**G**).

Given that complex neuronal circuits mediate social behaviors (Barak and Feng, 2016; Chen and Hong, 2018; Fernández et al., 2018), impairment of social novelty in *Fmr1*-KO mice may be due to abnormal neuronal activity in social-related brain regions, such as the anterior cingulate cortex (ACC) (Guo et al., 2019), hippocampus (Santini et al., 2013), and amygdala (Ferretti et al., 2019). We examined the expression of c-fos in these brain regions of WT and *Fmr1*-KO mice 1 hour after the social interaction **(Fig. 1E, F).** Strikingly, we detected fewer c-fos positive neurons in the RSG and ACC of *Fmr1*-KO mice **(Fig. 1E, G)**. The proportion of c-fos positive neurons in hippocampal CA1 and the basolateral amygdala did not change **(Fig. 1F, G)**. Previous studies improved that the retrosplenial granular cortex (RSG) has dense connections with social-related brain regions (Vogt, 2019a) and integrates different kinds of sensory information, including auditory, visual and olfactory (Jiang et al., 2018; Todd et al., 2016). These results raise our hypothesis that RSG may regulate social behavior. Also, since the RSG’s role in social interaction is unknown, we focused on the participation of the RSG in social interaction.

### Inactivation of RSG impairs social novelty behavior

To test our hypothesis, we examined the necessity of RSG to social interaction using the chemogenetic approach. We expressed the human M3 muscarinic DREADD receptor coupled to Gi (AAV-CamkIIα-hM4Di-mCherry) in CamkⅡα-positive neurons of RSG. With this approach, clozapine-n-oxide (CNO) could bind to the M3 muscarinic DREADD receptor, which activates the G protein inward-rectifying K^+^ channel and decreases the neuronal activity in the RSG. Two weeks after injection, the mice received systemic CNO (4.5 mg/kg, i.p, once per day) for one week before the three-chamber social interaction test (**Fig. 2A**). Both groups spent more time exploring the mouse in the sociability test than the empty cage (***fig. S2A & B***) 1 hour after CNO injection. In the social novelty test, the mice infected with hM4Di had no social preference for the unfamiliar mouse (**Fig. 2B & C**), which differed from those infected with the control virus. Therefore, the inactivation of RSG *via* the chemogenetic approach also impaired social novelty in mice. Similarly, we did not observe any movement-related changes in the open field (***fig. S2 C-E***).

**Figure 2.**
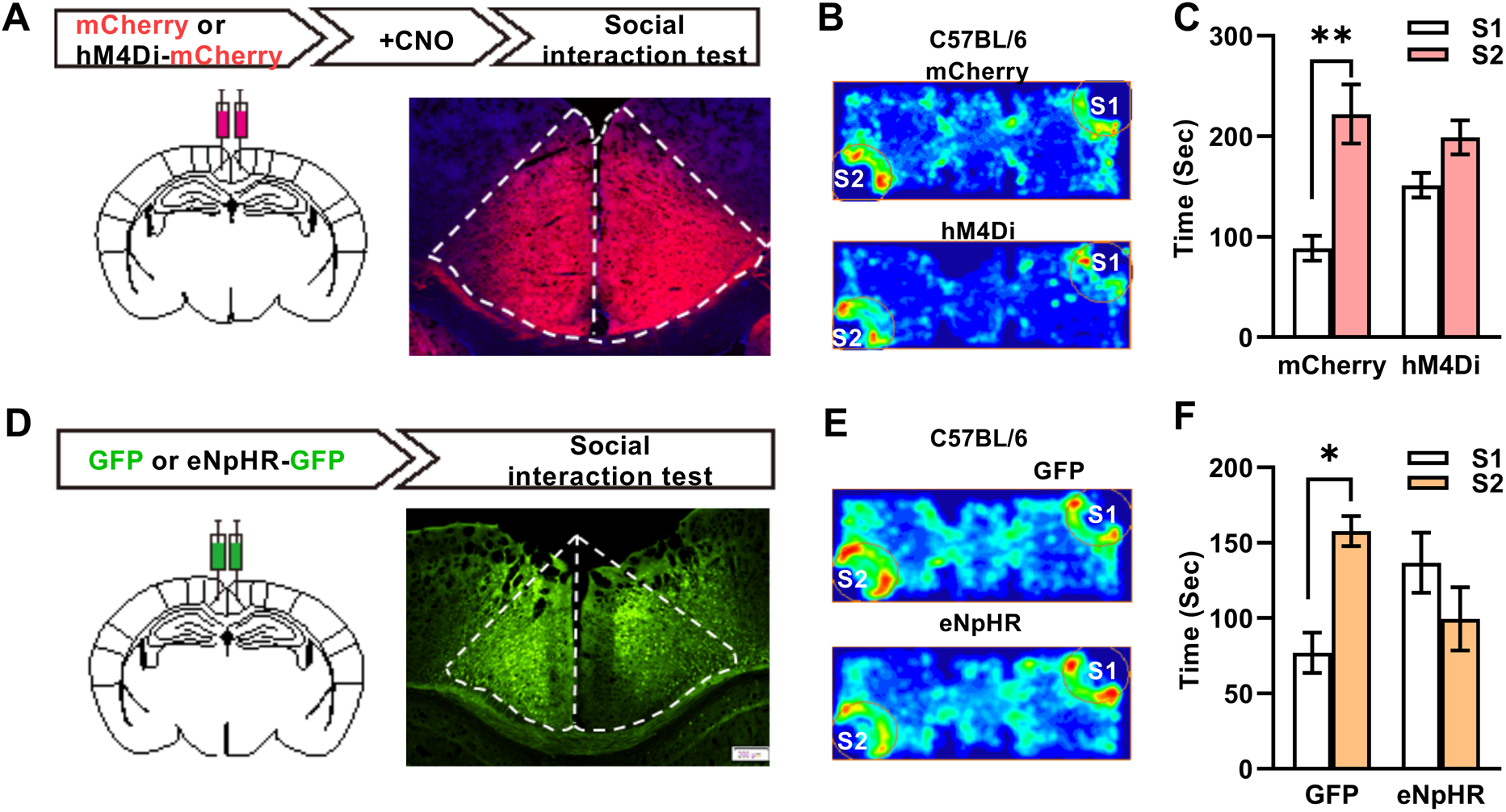
Inactivating RSG of normal mice impaired the social novelty. A, Upper: behavioral schedule and groups used; lower l left: brain diagram illustrating target areas manipulated; lower right: representative coronal section from an hM4Di-mCherry mouse. **B**, Representative mouse tracks heatmap during the social novelty test. **C**, Chemogenetic inhibition of RSG neurons impairs social novelty memory. Only the mCherry group prefer the novel over the familiar animal (mCherry, n = 6, *P* <0.01; hM4Di-mCherry, n = 10, *P* = 0.19, two-way ANOVA interaction, *F*_*(1,14)*_ = 3.83). RSG, retrosplenial granular cortex; ACC, anterior cingulate cortex; DG, dentate gyrus; BLA, basolateral amygdala. All heatmaps are scaled to the same color bar. Scale bar, 100 μm. **D**, Upper: behavioral schedule and groups used; lower left: diagram illustrating target areas manipulated; lower right: representative coronal section from an eNpHR-GFP mouse. **E**, Representative mouse tracks heatmap during the social novelty test. **F**, Light-inhibition of RSG neurons impairs social novelty memory. Only the GFP group prefer the novel over the familiar animal (GFP, n = 7, *P* <0.01; eNpHR-GFP, n = 6, *P* = 0.43, two-way ANOVA interaction, *F*_*(1,11)*_ = 8.13).

The chemogenetic approach may induce a long-term inactivation in RSG. We next examined the effects of transient inactivation of RSG on social interaction using the optogenetic approach. We expressed eNpHR in the RSG of C57BL/6 mice **(Fig. 2D)**. Two weeks after virus injection, A yellow laser was used to turn on eNpHR, a fast electrogenic Cl^−^ pump, which reduces the neural activity of viral labeled neurons (Gradinaru et al., 2008). We found that mice carrying eNpHR showed normal sociability (***fig. S2F & G***). However, in the social novelty test, mice with the yellow laser on spent the same amount of time interacting with the familiar and the novel mice **(Fig. 2E, F)**. On the contrary, the laser did not affect mice’s performance with the control virial (AAV2/8-CamkⅡα-eGFP) in the social novelty test. These mice with eNpHR had normal anxiety level (***fig. S2H-J***), indicated by similar exploration time in the central region and running distance of the open field. Further experiments confirmed the expression of eNpHR in the RSG (**Fig. 2D**). Therefore, transient inactivation of RSG also impairs social novelty, but not sociability. In summary, both inhibition of RSG excitatory neurons transient and long-termly impaired the social interaction of C57BL/6 mice to novelty social objects.

### Activation of RSG rescues the social novelty deficit in*Fmr1*-KO mice

Based on the social novelty deficits during RSG inactivation and the low c-fos expression in RSG of *Fmr1*-KO mice after the social interaction test, we hypothesized that social novelty deficit in *Fmr1*-KO mice might be due to the low neural activities in RSG. To test this hypothesis, we optogenetically activated the excitatory neurons in RSG of Fmr1-KO mice. We expressed ChR2 under the premotor of CamkⅡα in RSG of both hemishperes and implanted an optical fiber 0.3 mm above the virus injection sites 18 days later (**Fig. 3A**). To validate the excitatory effect of ChR2 on RSG neurons, we performed whole-cell patch-clamp recordings in ChR2-expressing neurons and found marked bursts of action potentials under blue laser light stimulation (**Fig. 3B**). One week after optical fiber implantation, the mice have habituated the environments for two days (see methods). Then, social behavior was tested in *Fmr1*-KO mice during which excitatory RSG neurons were activated by blue laser light (20 Hz, 10 mW). The mice with ChR2-eGFP or eGFP only behaved as usual in the sociability test (***fig. S2A &B***). Strikingly, the *Fmr1*-KO mice with ChR2 expression showed typical WT like social novelty responses, in that they presented apparent social preference for the unfamiliar mouse (**Fig. 3C, D**). In contrast, the *Fmr1*-KO mice expressing control virus (eGFP) showed an inadequate social novelty response (**Fig. 3C, D**). These data suggested that transient increasing the activity of excitatory RSG neurons is sufficient to rescue the social novelty deficits in *Fmr1*-KO mice.

**Figure 3.**
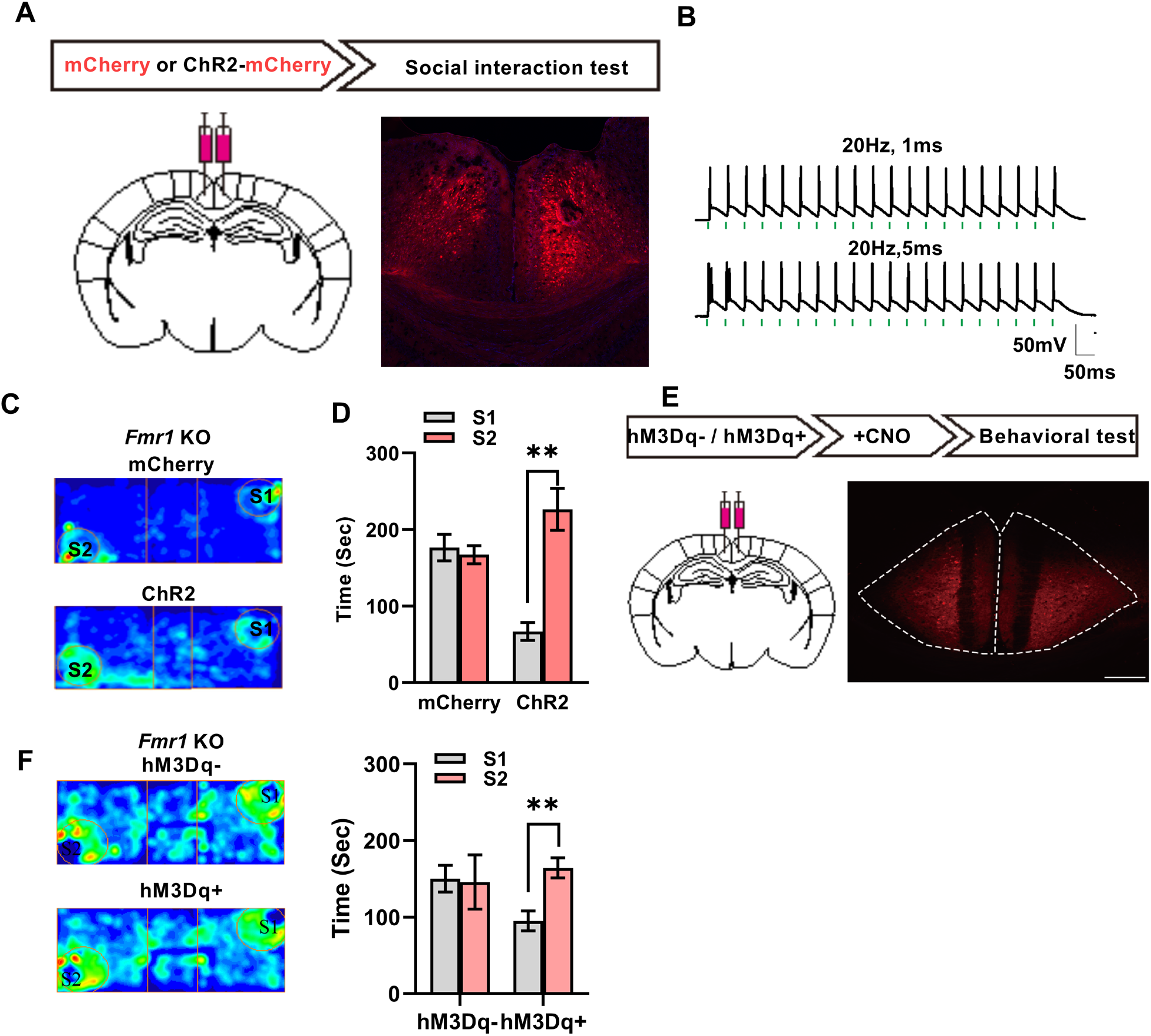
Optogenetic and chemogenetic activation of the RSG rescues the response to social novelty in Fmr1-KO mice. **A**, Upper: behavioral schedule and groups; lower left: diagram illustrating target areas manipulated; lower right: representative coronal section from a ChR2-mCherry mouse. **B**, Action currents recorded from RSG pyramidal neurons expressing ChR2-mCherry in response to 20, 20 Hz, 1-ms or 5-ms blue (470 nm) light pulses. **C**, Representative mouse tracks heatmap during the social novelty test. **D**, Light-activation of RSG neurons rescues the response to social novelty in Fmr1-KO mice. The ChR2-mCherry rescue group prefer the novel over the familiar animal (mCherry, n = 10, *P* = 0.90; ChR2-mCherry, n = 5, *P* <0.01; two-way ANOVA interaction, *F*_*(1,13)*_ = 18.30). **E**, Upper: behavioral schedule and groups; lower left: diagram illustrating target areas manipulated; lower right: representative coronal section from an hM3Dq-mCherry mouse. **F**, Representative mouse tracks heatmap during the social novelty test. **G**, Chemogenetic activation of RSG neurons rescues the response to social novelty in Fmr1-KO mice. The hM3Dq+ rescue group prefer the novel over the familiar animal (hM3Dq−, n = 3, *P* = 0.98; hM3Dq+, n = 6, *P* <0.01; two-way ANOVA interaction, *F*_*(1,7)*_ = 6.64). All heatmaps are scaled to the same color bar. Scale bar, 200 μm.

To further confirm this point, we used a chemogenetic approach by expressing the human M3 muscarinic DREADD receptor coupled with Gq (hM3Dq) in CamkIIα-positive neurons in the RSG of *Fmr1*-KO mice. Binding of CNO to this receptor increased neural activity. After the stable expression of hM3Dq (**Fig. 3E**), we also systemically applied CNO (4.5 mg/kg, once per day for one week) before the three-chamber social interaction test. The CNO application did not change sociability (***fig. S3 C & D***). In the social novelty test, *Fmr1*-KO mice with hM3Dq spent more time with the unfamiliar mouse, while *Fmr1*-KO mice with control virus did not (**Fig. 3F & G**). Our chemogenetic data confirmed the optogenetic results that increasing neural activity of the RSG long-termly is sufficient to rescue the social novelty deficits in *Fmr1*-KO mice.

### Social interaction activates fewer neurons in the RSG of*Fmr1*-KO mice

All activation results in *Fmr1*-KO mice and inactivation results in WT mice indicated that RSG is a crucial area for social novelty but does not influence sociability. We hypothesized that neural ensemble activity in the RSG from the WT and KO mice might differ during social interaction behaviors. To test this hypothesis, we expressed GCaMP6s, a fluorescent Ca^2+^ indicator, in the RSG *via* the AAV virus. Then, with a miniature fluorescent microscope mounted on the mouse’s head, we imaged neuronal activity in the RSG during the three-chamber sociability and social novelty tests (**Fig.4A & *Fig. S4A***). The WT mice showed normal sociability and social novelty, while *Fmr1*-KO mice showed an impaired social novelty response. These results indicated no social behavior deficits when RSG neuronal activity was imaging with a head-attached miniature microscope (**Fig.4 A & B**). We then analyzed neuronal activity in the RSG when mice performed the three-chamber social task (**Fig.4C**). We found that *Fmr1*-KO animals showed higher neuronal activity than the controls during social interaction tests (***Fig. S4***), indicating a difference in the background neuronal activity in *Fmr1*-KO mice.

**Figure 4.**
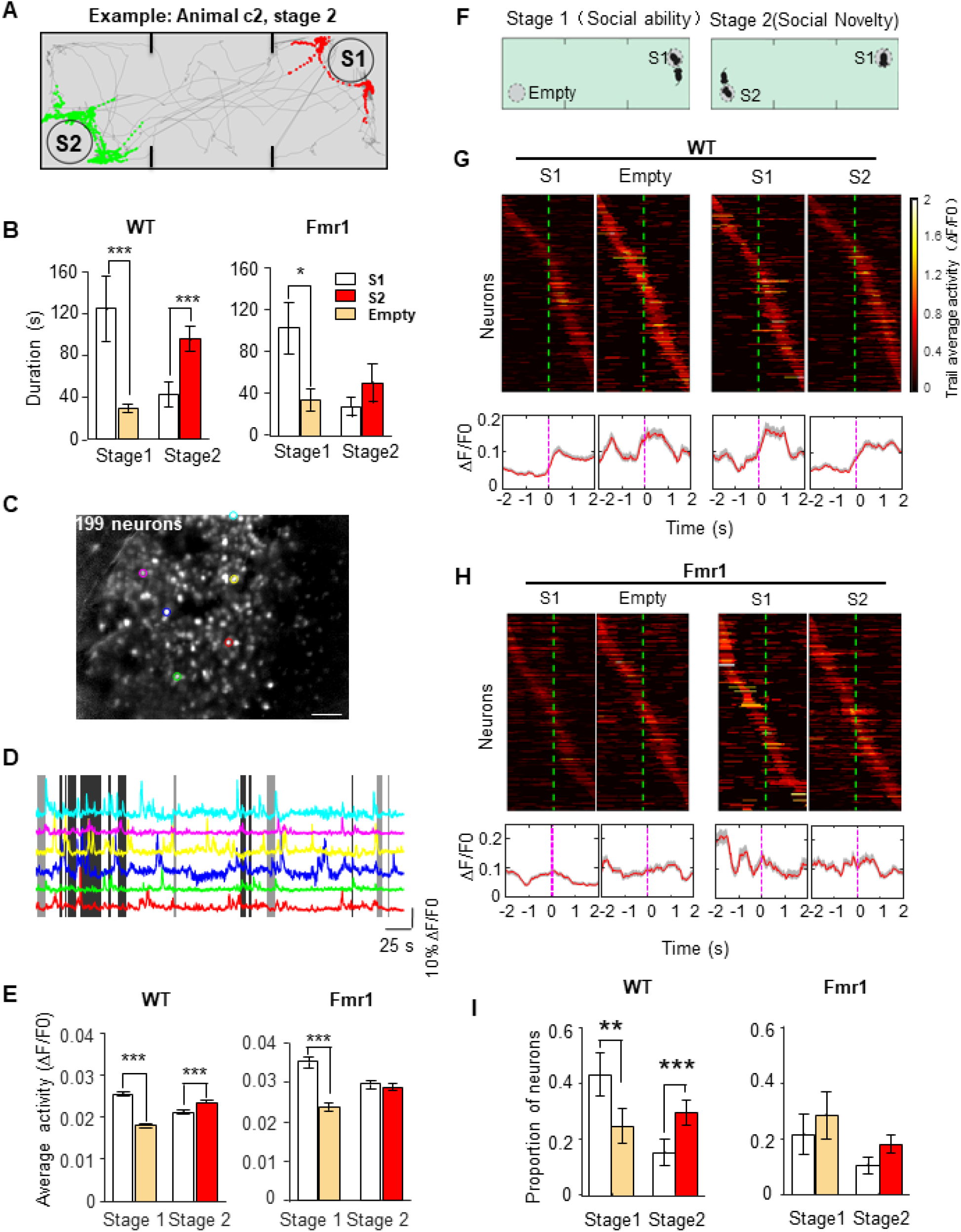
Social interaction activates fewer neurons in the RSG of *Fmr1*-KO mice. **A**, Example movement trace during the social interaction test. Gray line, movement trace; red dots, social interaction with a familiar animal (**S1**); green dots, social interaction with an unfamiliar mouse. **B**, *Fmr1* KO mice show impaired social novelty during Ca*2+* imaging. Both WT and KO mice prefer Strange 1 over an empty chamber (WT, n = 6 mice, *P* <0.001; KO, n = 4, *P* < 0.05). Only WT mice prefer the novel mouse (S2) over the familiar mouse (WT, *P* <0.001; KO, *P* >0.05). **C**, Representative standard deviation projection of 15,000 ΔF/F image frames collected over 7.5 min in the retrosplenial cortex during the social novelty test paradigm. White dots indicate identified neurons, and 199 neurons are identified from this animal. Scale bar, 100 um. **D,** Example Ca^2+^ traces from ROIs with matched colors in C. Gray areas, Ca^2+^ transients during social interaction with strange animals; black areas, with familiar animals. **E**, *Fmr1* KO mice show less neuronal activity during the social novelty test. We calculated the integrated Ca^2+^ activity during the social interaction period. Both WT and KO mice RSG neurons showed more activity during interaction with the Strange 1 over an empty chamber (WT, n = 6 mice and 721 neurons, *P* <0.001; KO, n = 4 animals, and 390 neurons, *P* <0.001). Only neurons in WT mice showed more activity during interaction with the novel Strange 2 mouse over the familiar mouse (WT, n = 6 mice and 679 neurons, *P* <0.001 KO, n = 4 mice and 360 neurons. *P* = 0.3). **F**, Left: social ability test. Right: social novelty test. **G,** Trial-average activity of retrosplenial cortex neurons in WT animals showing increasing activity when interaction begins. The active event traces from different trials are aligned to the initial contact with different objects (vertical dash lines), then averaged across trials. All the social interaction-related neuronal activity traces are sorted based on their peak activation time and displayed in a temporal raster plot. Neurons with no activity in the task were removed. The left two columns indicate the activity in stage 1, and the right two columns indicate the activity in stage 2. **H,** Trial-average neuronal activity in Fmr1-knockout mice. **I,** Proportions of social-interaction neurons.

To further dissect the neural ensemble activity during social interactions, we first analyzed the calcium events of imaging animals during direct contact with the targets. We found that WT animals showed stronger fluorescence intensity when they contacted animal than empty cups in the social ability test (stage 1), and higher activity in contacting novel animal than the familiar mouse in the social novelty test (**Fig.4 D & E,** stage 2). Unlike the WT mice, neuronal activities were higher when the *Fmr1*-KO mice contacted S1 than empty cups in the sociability test. However, it was similar when KO mice contacted novel animals and familiar animals in the social novelty test (**Fig.4 D & E**, stage 2). Some evidence indicates that neural activity in the PFC increases when animals begin to contact other animals (Barbera et al., 2016). To test whether similar effects occur in the RSG, we aligned the neuronal activity to the time point when test animals contact targets (S1, S2, and empty cups) (**Fig.4 F - H**). We found higher neuronal activity in the RSG when WT animals began to contact all targets. In contrast, there were no changes in the *Fmr1*-KO animals (**Fig.4F - H**).

Furthermore, we found that more neurons were activated when WT mice contacted S1 than empty cups in the sociability test (stage 1) and more neurons in novel animals than the familiar animal in the social novelty test **(Fig.4I)**. However, *Fmr1*-KO mice showed similarly activated neurons to all targets (S1 and empty cup in stage1, S1, and S2 in stage 2) **(Fig.4I).** So, the activity of the RSG during social interactions correlated with the social behaviors. These results indicated that social novelty increases the neuronal activity in the RSG of WT mice, but not in *Fmr1*-KO mice.

### Social experience input to RSG in the sociability test stage is necessary for the social novelty response

The manipulation of RSG only impaired the social novelty, but not in sociability. One possibility is that RSG neurons receive and store social-related information about the S1 animal in stage 1, which is necessary for the social behaviors in the social novelty. If this is right, inhibiting neuronal activity in the RSG during sociability will impair the social novelty response. To test this hypothesis, we also performed the optogenetic inactivation in sociability test. Similarly, we expressed the eNpHR in the excitatory neurons of RSG and implanted the optical fibers. One week after fiber implantation, we performed a social interaction test and applied yellow laser light at stage1 (**Fig. 5A**). Similarly, inhibiting the RSG did not change sociability (**Fig. 5B**). However, the mice with eNpHR spent equal amounts of time exploring the familiar and unfamiliar mice in social novelty. While the yellow laser did not affect mice with control virus (eGFP) (**Fig. 5 C & D**). This result suggested that RSG neurons do not regulate sociability. However, the neuronal activity in the sociability test is necessary for the standard social novelty behavior.

**Figure 5.**
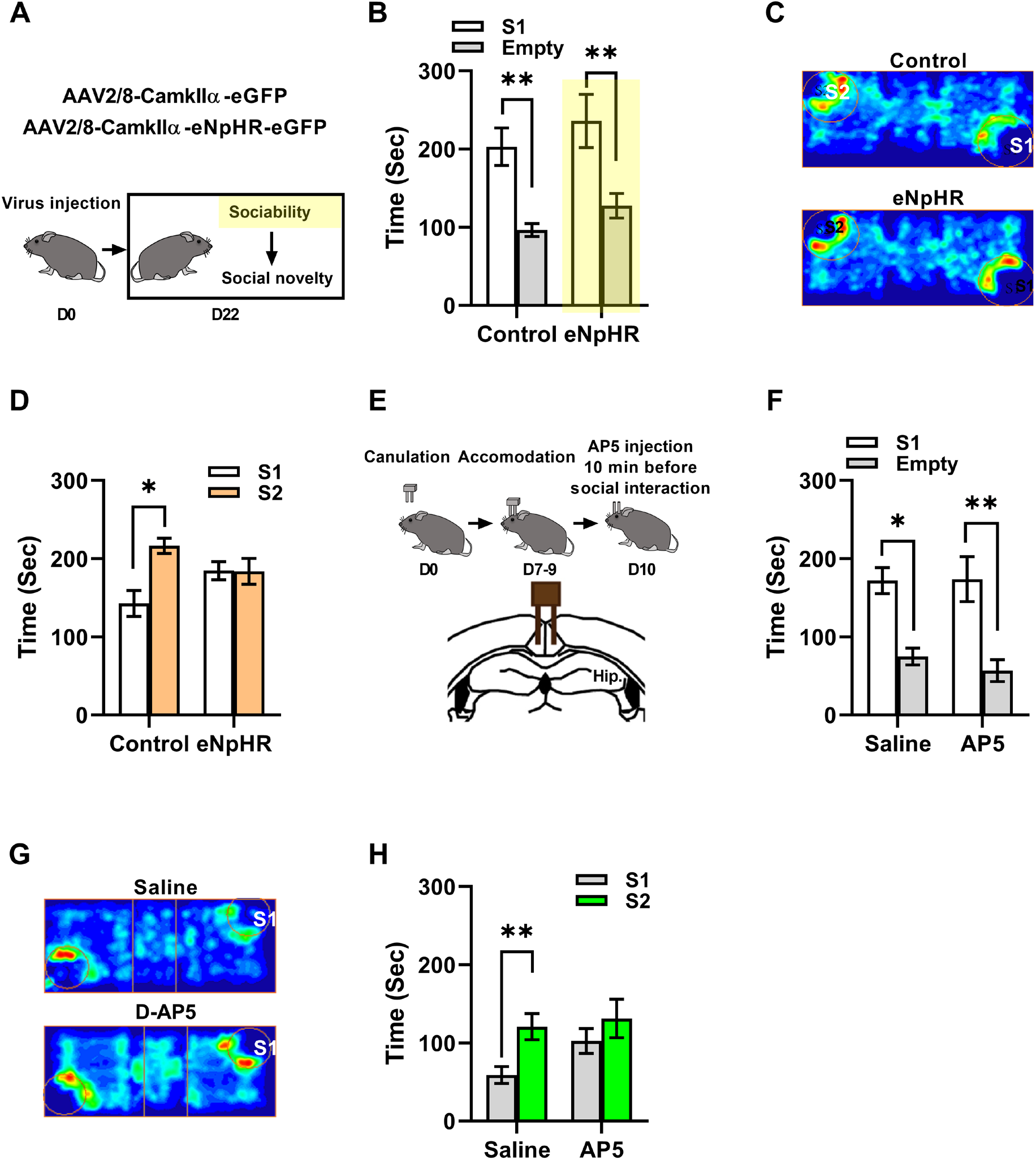
Inputs to the RSG during the sociability test are necessary for the response to social novelty. **A**, Protocol for optogenetic inhibition during the sociability test stage. **B**, Both the control and eNpHR groups show intact sociability (Control, n = 7, *P* <0.01; eNpHR, n = 7, *P* <0.01; two-way ANOVA interaction, *F*_*(1,12)*_ = 0.002). **C**, Representative mouse tracks heatmap during the social novelty test. **D**, Only the control group shows intact social novelty (Control, n = 7, *P* <0.05; eNpHR, n = 7, *P* = 0.99; two-way ANOVA interaction, *F*_*(1,12)*_ = 5.14). **E**, Upper: protocol for AP5 injection 10 min before the sociability test stage; lower: diagram illustrating target areas for cannula implants. **F**, Both the saline and AP5 groups show intact sociability (Saline, n = 7, *P* <0.05; AP5, n = 11, *P* <0.01; two-way ANOVA interaction, *F*_*(1,16)*_ = 0.18). **G**, Representative mouse tracks heatmap during the social novelty test. **H**, Only the saline group shows intact social novelty (Saline, n = 7, *P* <0.01; AP5, n = 11, *P* = 0.13; two-way ANOVA interaction, *F*_*(1,16)*_ = 2.00). All heatmaps are scaled to the same color bar.

N-methyl-D-aspartate receptor (NMDAR) is one of the ionotropic glutamate receptors in the cerebral cortex, critical for higher cognitive functions. The restoration of NMDAR functions rescues social behaviors in the Shank2-KO mouse, a model of autism (Won et al., 2012). In the RSG, the input information would activate NMDARs. We further examined the role of NMDARs in mediating synaptic transmission during social interaction by micro-infusion of DL-AP5, a selective antagonist of NMDARs. Mice received adequate accommodation one week after cannula implantation, and we then ran the social interaction test after bilateral micro-infusion of DL-AP5 (**Fig. 5E**). While blocking the activity of NMDARs did not affect the sociability test (**Fig. 5F**), mice treated with DL-AP5 exhibited an impaired social novelty response, but not those with the vehicle (**Fig. 5G & H**). Therefore, the NMDARs mediating information input in the RSG are necessary for a social novelty response.

### The innervations from the subiculum (SUB) to the RSGs contributes to social interaction

We next investigated the possible pathways that convey the social experiences to the RSGs by injecting the AAV2/retro-CamkⅡα-EGFP into the RSGs (**Fig. 6A**). Interestingly, we observed the sparsely labeled neurons in the ACC, but densely labeled neurons in the subiculum (SUB) and the anteroventral thalamic nucleus (AV) three weeks after the virus injection (**Fig.6A**), which indicates that the RSG receives the innervations from these brain regions. To selectively manipulate the related pathways, we injected the pAOV2/retro-CamkⅡα-eYFP-2A-Cre into the RSG first (D0), and injected the AAV-Ef1α-DIO-eNpHR3.0-mCherry/AAV-Ef1α-mCherry into the ACC or SUB 10 days later, separately. The expressing of Cre in the ACC or SUB will perform recombination on the sequence between LoxP sites, therefore induced the expression of eNpHR3.0 in the projection neurons in ACC or SUB; therefore, we can selectively manipulate the ACC-RSG or SUB-RSG pathway by deliverer the yellow laser (**Fig. 6B**). Optogenetically inactivation of the ACC-RSG pathway impaired the social ability (**Fig. 6C, D**), but not the social novelty (**Fig. 6E**). Using the same approach, we inactivated the SUB-RSG pathway (**Fig. 6F**), which also impaired sociability (**Fig. 6G, H**). However, the mice with eNpHR3.0 showed altered social novelty behaviors; they spent less time sniffing the unfamiliar mouse than the Ctrl group (**Fig. 6I**). These data suggest that the SUB-RSG neurons are critical for sociability and may send social-related information to the RSG for the social novelty test.

**Figure 6.**
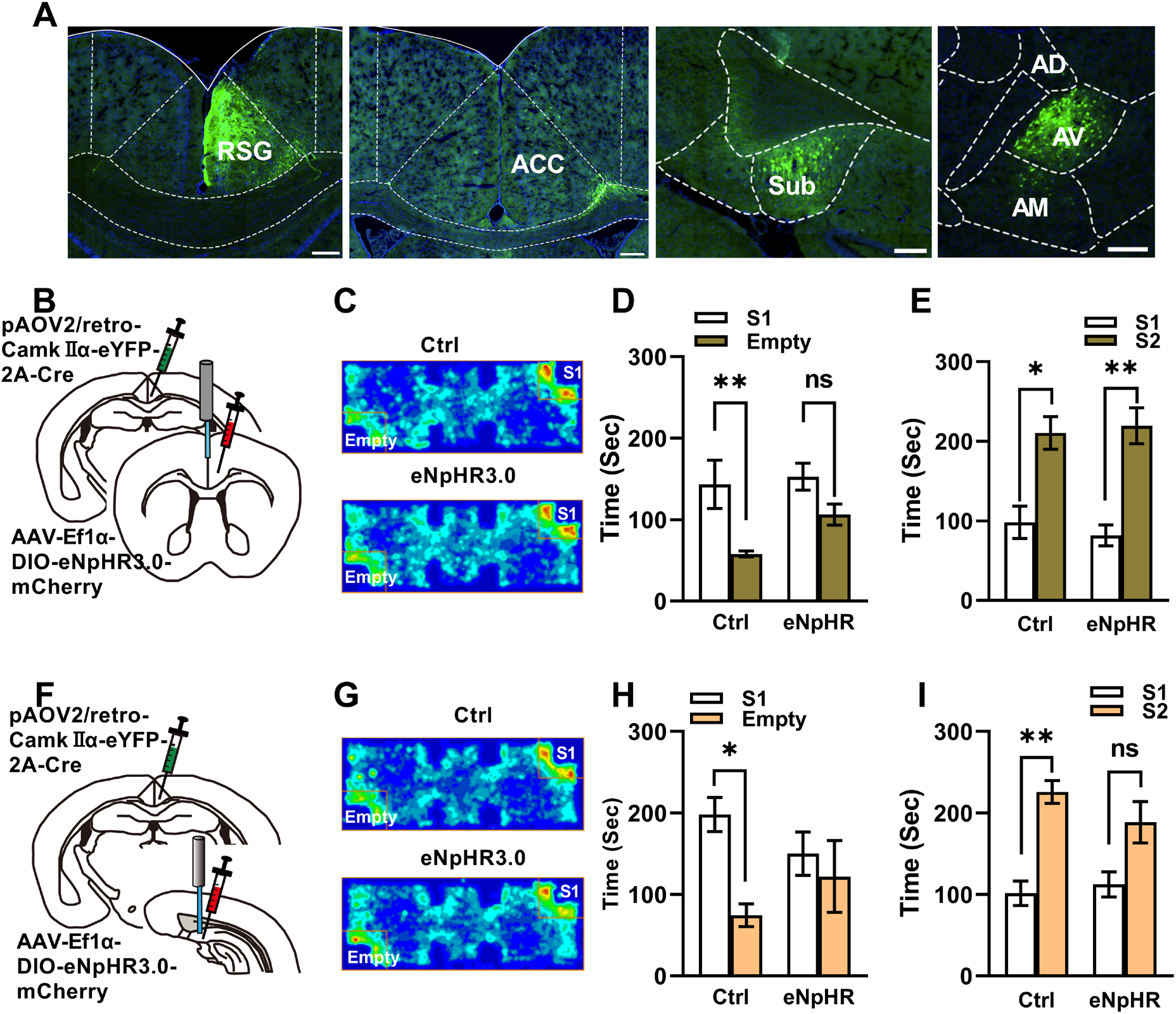
The innervations from the subiculum (SUB) and ACC contributes to the social interactions. A. The RSG receives the innervations from the ACC, SUB and anteroventral thalamic nuclus (AV). B. Diagram illustrating target area for cannula implants and virus injection to examine the role of ACC-RSG pathway in social interaction. C. Representative mouse tracks heatmap during the sociability test with ACC-RSG pathway interaction. D. Inactivation of ACC-RSG pathway impaired sociability (Ctrl, n = 5, *P* <0.01; eNpHR, n = 6, *P* = 0.09; two-way ANOVA interaction, *F*_*(1,9)*_ = 1.75). E. Inactivation of ACC-RSG pathway did not affect social novelty (Ctrl, n = 5, *P* <0.05; eNpHR, n = 6, *P* < 0.01; two-way ANOVA interaction, *F*_*(1,9)*_ = 0.29). F. Diagram illustrating target area for cannula implants and virus injection to examine the role of SUB-RSG pathway in social interaction. G. Representative mouse tracks heatmap during the sociability test with SUB-RSG pathway inactivation. H. Inactivation of SUB-RSG pathway impaired sociability (Ctrl, n = 6, *P* <0.05; eNpHR, n = 6, *P* = 0.50; two-way ANOVA interaction, *F*_*(1,10)*_ = 1.75). I. Inactivation of ACC-RSG pathway also affected social novelty (Ctrl, n = 6, *P* <0.01; eNpHR, n = 6, *P* = 0.09; two-way ANOVA interaction, *F*_*(1,10)*_ = 1.04).

## Discussion

In the current study, we investigated the roles of the RSG in social interaction by combining multiple approaches and found that neuronal activity in the RSG was necessary for social novelty but not sociability during the social interaction test. Decreasing RSG neuronal activity impaired the response to social novelty in normal mice, and enhancing this activity rescued the abnormal response in *Fmr1*-KO mice. Moreover, the RSG roles in social novelty are dependent on the information input in the sociability test. Our results strongly suggest that the RSG is one of the brain areas which is essential for social cognition. This helps to understand the neural circuits of social awareness and provides a potential method to treat Autism spectrum disorders in the future.

The RSG is part of the complex neural circuitry that mediates social cognition (Chen and Hong, 2018). It has dense connections with the brain regions related to social cognition at the neuronal circuit level; it connects to the auditory cortex, visual cortex, PFC, thalamus, and hippocampus in humans, rats, and macaque monkeys (Barrière et al., 2019; Rolls, 2019; Yamawaki et al., 2019). Different brain regions regulate social behaviors, receiving social clues, comparing them with stored information, making decisions, and performing social interactions (Chen and Hong, 2018). The RSG may receive and integrate different sensory information directly from the auditory cortex, visual cortex, thalamus, and PFC to achieve social interaction (Rolls, 2019; Vogt, 2019b).

Previous studies have shown that the RSG is involved in episodic memory, navigation, imagination, and planning for the future (Vann et al., 2009). Here we found fewer c-fos positive neurons in the RSG of *Fmr1*-KO mice after social interaction. Moreover, increasing the activity of RSG neurons *via* optogenetic or chemogenetic approaches rescued the response to social novelty in *Fmr1*-KO mice. Combined with the results from WT mice, our data strongly suggest that the RSG is essential for the response to social novelty. Considering its tight connections with the hippocampus (Yamawaki et al., 2019), the RSG may also process spatially-related social information.

Although inactivating the RSG did not affect sociability, our Ca^2+^ imaging results clearly showed that activating RSG neurons during this period and blocking the activity of NMNDARs with DL-AP5, specifically in the RSG, impaired the response to social novelty. Therefore, the information received by the RSG in stage one is critical for social novelty. Here, we propose that the RSG is one of the brain regions that change the internal state based on information received at stage1 and perform social behaviors at the stage2 (Chen and Hong, 2018). Using the retrograde tracing approach, we identified three brain regions, including the ACC, SUB, and AV of the thalamus, that may send the social-related information to the RSG. The selective inactivation of the SUB-RSG pathway impaired both sociability and social novelty, indicating that the SUB-RSG pathway may convey the social experience to the RSG, contributing to the social novelty. The ACC-RSG may also contribute to the sociability; while the inactivation did not affect the social novelty, the RSG may receive social-related information from other pathways. Due to the limitation, here we just identified three regions innervates to the RSG, there must be some other sources that innervate to the RSG, and other tracing approaches should be used for further studies.

Modulation of the neuronal activity in social-related brain regions may rescue the social deficit in FXS. A pharmaceutical approach targeting mGluR and GABAb receptors have shown less efficiency in clinical trials (Berry-Kravis et al., 2018). In the current study, we showed that activating the RSG can change social behaviors, providing a strategy that enhanced RSG neuronal activity in patients with FXS could rescue the social deficits. Repetitive transcranial magnetic stimulation is a form of noninvasive brain stimulation used to manage autism and FXS (Oberman et al., 2010, 2016), such stimulation of the RSG may be an effective and safe approach to managing FXS in the future.

## Acknowledgments

The authors would like to thank Hailan Hu, Junyu Xu, and Ji Hu for their critical comments on this manuscript. We thank Dr. San-Hua Fang at the Core Facilities of the Zhejiang University School of Medicine for technical assistance. The authors apologize to colleagues whose work could not be cited due to space and reference restrictions.

## Author contributions

SHF performed behavioral tests, western blots, immunostaining, and data analysis; CRL, SH and LXJ performed *in vivo* Ca^2+^ imaging and analyzed the data; LYN made the whole-cell patch-clamp recordings; LL, CW, and GLX performed the behavioral tests and analyzed the data, CZ provided the *Fmr1* KO mice and analyzed data; LJH, LXJ, and LXY designed the experiments and approved the draft; and LXY wrote the paper.

