## Supplemental figure 1-4 for "Deconstruction of the retrosplenial granular cortex for social behavior in the mouse model of fragile X syndrome"

Figure S1-S4

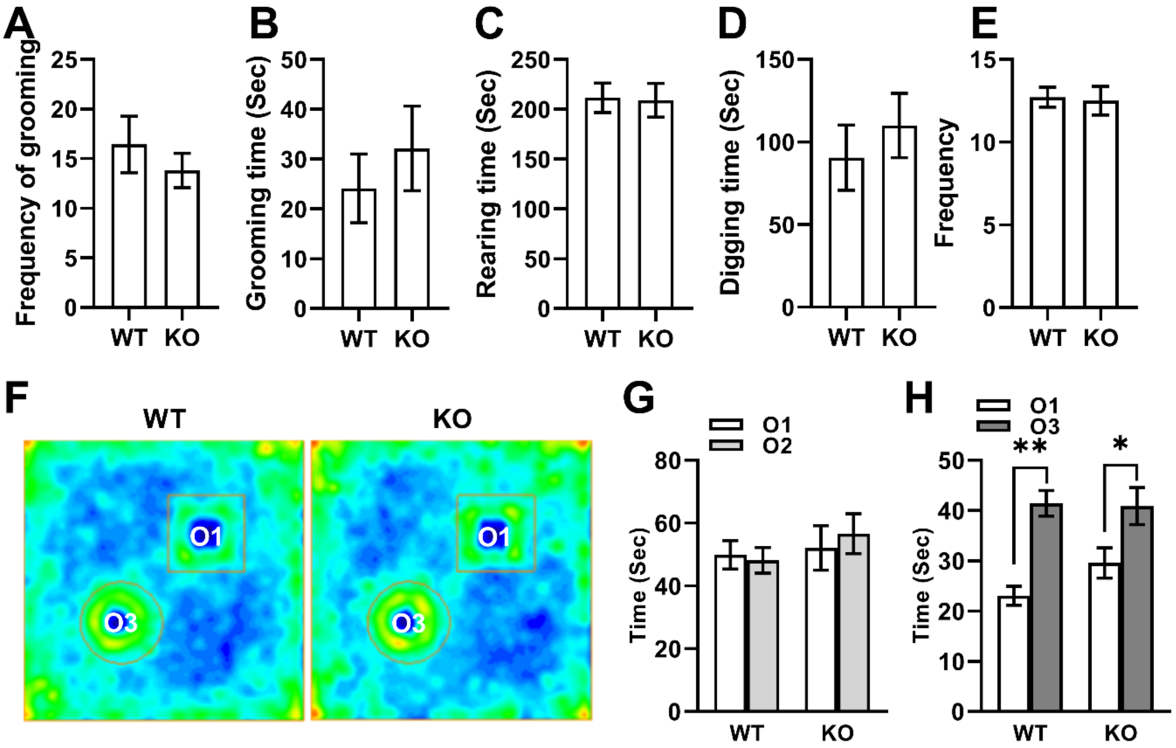


**Figure S1. *Fmr1* KO mice showed normal stereotypical behaviors.**

**A**, no significant difference between WT (n = 7) and KO (n = 5) mice in frequency of grooming (*P* = 0.48).

**B**, No significant difference between WT and KO mice in grooming time within five minutes (*P* = 0.91).

**C**, no significant difference between WT and KO mice in rearing time (*P* = 0.91).

**D**, no significant difference between WT and KO mice in digging time ($P = 0.51$).

**E**, no significant difference between WT and KO mice in the frequency of digging ($P = 0.84$).

F. Representative mouse tracks heatmap during the stage2 of the novel objects recognition test (NOR).

G. Both the WT and KO mice spent equal time on the two identical objects in stage1. (WT, n = 15, KO, n = 13).

H. Both the WT and KO mice spent longer time on the novel object in stage2 of NOR test. (two-way ANOVA, interaction, $F_{(1,25)} = 2.29$, $P > 0.05$; WT vs. KO: $F_{(1,25)} = 0.90$, $P = 0.35$; objects: $F_{(1,25)} = 40.08$, $P < 0.01$;WT, n = 15, KO, n = 13).

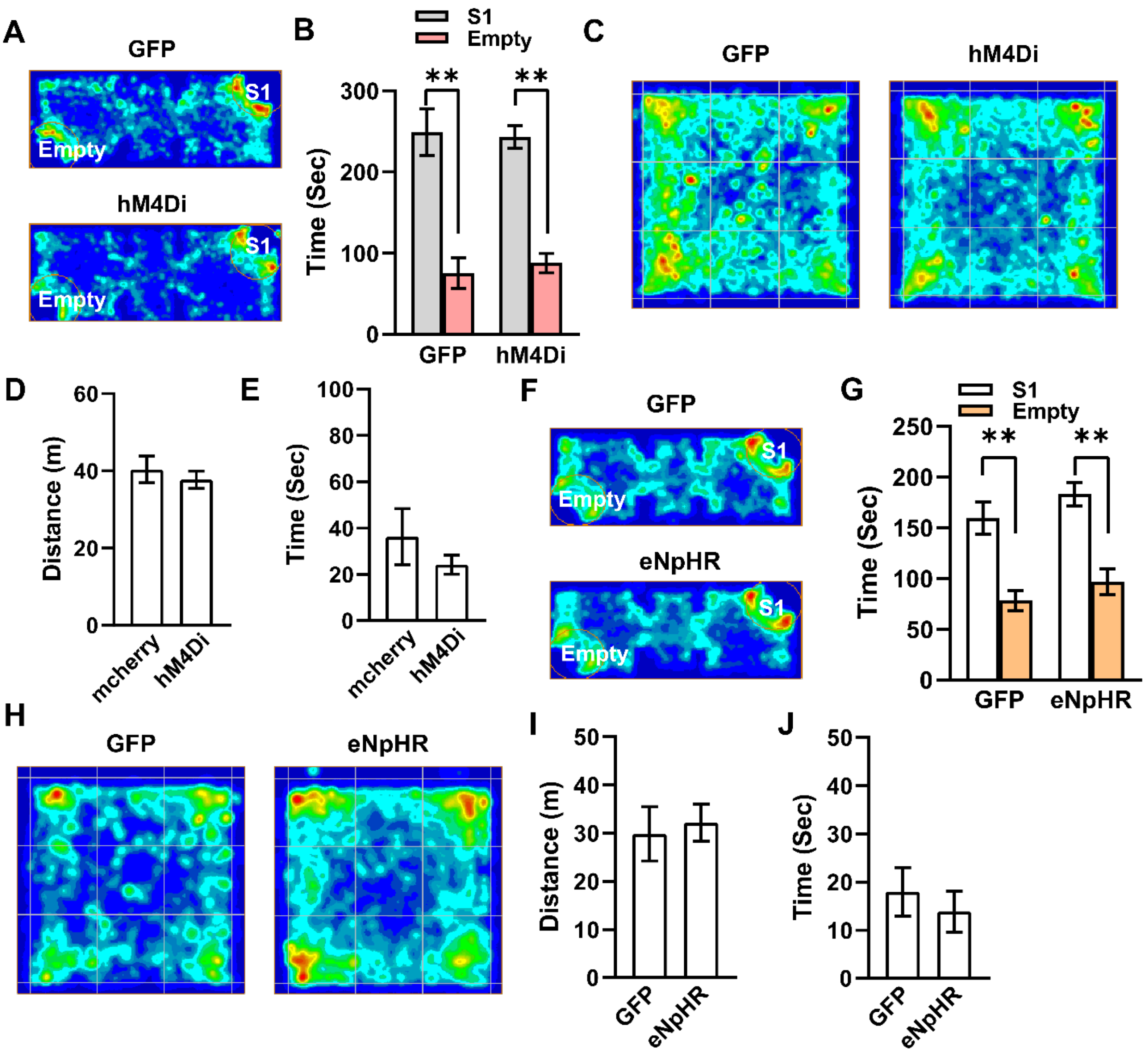


**Figure S2. Inactivating RSG by optogenetic and chemogenetic approaches showed normal sociability and Locomotor Activity.**

**A**, Representative mouse tracks heatmap during sociability test.

**B**, Both GFP and hM4Di group mice showed the intact sociability (GFP, n = 6, $P < 0.01$; hM4Di, n = 10, $P < 0.01$. Two-way ANOVA, $F_{(1, 28)} = 0.27$).

**C**, Representative mouse tracks heatmap during the open field test.

**D**, no significant different between GFP and hM4Di group mice in total traveling distance (GFP, n=6, hM4Di, n =7, unpaired t-test, $P = 0.53$).

**E**, no significant different between GFP and hM4Di group mice in total travelling time (GFP, n=6; hM4Di, n =7; unpaired t-test, $P = 0.33$). All heatmap shown is scaled to the same color bar.

**F**, Representative mouse tracks heatmap during sociability test.

**G**, Both GFP and eNpHR group mice showed the intact sociability (GFP, n = 7, $P < 0.01$; eNpHR, n = 6, $P < 0.01$; Two-way ANOVA, $F_{(1, 22)} = 0.04$).

**H**, Representative mouse tracks heatmap during the open field test.

**I**, no significant difference between GFP and eNpHR group mice in total traveling distance (unpaired t-test, $P = 0.73$).

**J**, no significant difference between GFP and eNpHR group mice in total traveling time (unpaired t-test, $P$ =0.54).

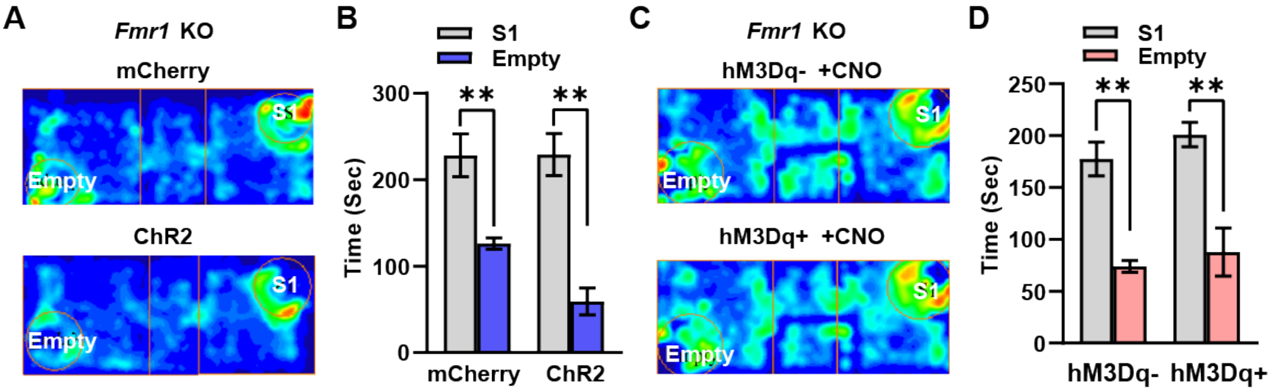


**Figure S3. Activating RSG by optogenetic and chemogenetic approaches showed normal sociability.**

**A**, Representative mouse tracks heatmap during sociability test.

**B**, Both GFP and ChR2 group mice showed the intact sociability (GFP, n = 10, $P$ <0.01; ChR2, n = 5, $P$ <0.01; two-way ANOVA, $F_{(1, 26)}$ = 2.631).

**C**, Representative mouse tracks heatmap during sociability test.

**D**, Both GFP and hM3Dq group mice showed the intact sociability (GFP, n =3, $P$ =0.01; hM3Dq, n = 6, $P$ < 0.01; two-way ANOVA interaction, $F_{(1, 14)}$ = 0.06 ). All heatmap shown are scaled to the same color bar.

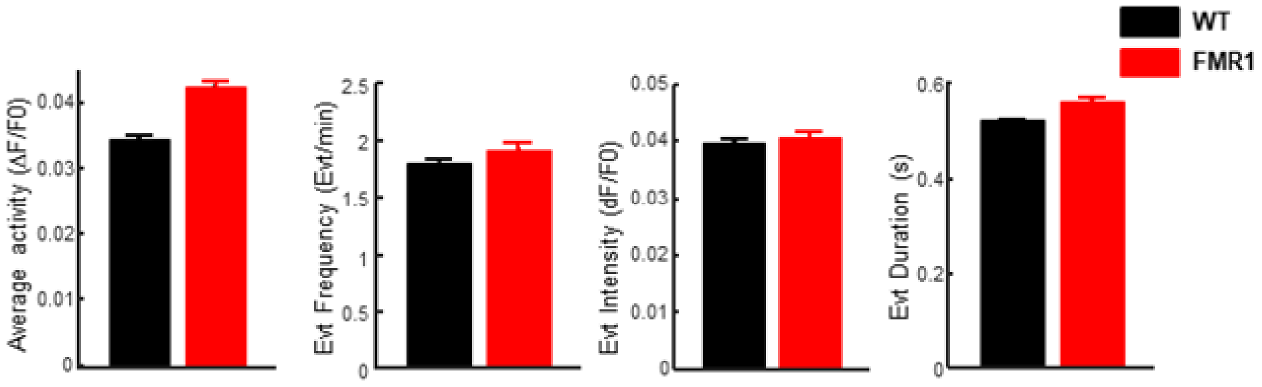


**Figure S4.** ***Fmr1*** **KO animals showed higher neuronal activities than the controls during social interaction tests.** Quantifications of identified integrated calcium transient activity, calcium transient frequency, calcium transient amplitude, and calcium transient frequency duration of WT and Fmr1 animals. Error bars representing SEM.
